# Simulation of enteric colonisation by and screening for Carbapenemase Producing Enterobacteriaceae using an in-vitro human gut model

**DOI:** 10.1101/399170

## Introduction

Carbapenemase Producing Enterobacteriaceae (CPE) are increasing worldwide [1, 2] and pose a significant threat to public health. The burden of CPE infection is multifaceted, encompassing adjustment of treatment regimens [3, 4], increased duration of inpatient stay [5], associated morbidity [6] and mortality [5, 7, 8], in addition to the wider burden it poses on healthcare systems, with financial and societal implications [9]. The rapid spread of CPE within endemic areas [10] and in the health care setting [11] is well documented. Furthermore, illustrates the capacity of CPE exposure to lead to gut colonisation [12].

Sensitive screening methods are needed to identify CPE infection or colonisation, to ensure adequate infection control mechanisms can be instigated as soon as possible. Public health prevention strategies have proven vital to the control of CPE in previous endemic areas [13] and healthcare based outbreaks [14]. Once an individual has been exposed to CPE little is known regarding the colonisation process, including the infective dose and duration of exposure required for colonisation, disruption to the normal gut flora and the impact of antibiotic administration [15, 16]. The choice and duration of antimicrobial therapy can pose a significant clinical dilemma in patients that have a positive screening test

We have used a well validated clinically reflective *in-vitro* human gut model [17] to evaluate common screening methods for the detection of CPE and to follow the growth of CPE populations within the human gut microbiota. The selected CPE strains inoculated into the model represent those of major clinical significance; comprising of strains encoding KPC, OXA 48, NDM genes [18]. Five different screening tests were used, comprising both selective media and molecular methods.

## Methods

### In vitro gut model

The gut model utilised in this study is based on that of MacFarlane *et al*, and has been validated against physico-chemical and microbiological properties of the colonic contents of sudden death victims [17]. It is a triple-stage continuous culture system, arranged in a weir cascade formation, that simulates the proximal, medial and distal sections of the human colon. The three vessels are maintained at 37°C (via circulating waterbath), at physiologically relevant pH and volume (Vessel 1 - 280mL, pH 5.5±0.2; Vessel 2 - 300mL, pH 6.2±0.2; Vessel 3 - 300mL, pH 6.8±0.2). The system is continuously sparged with nitrogen to maintain anaerobicity, and top fed with a complex growth media [19] at a controlled rate (D=0.015 h-1; system retention time 66.7h). The gut model is inoculated with pooled faecal slurry (10% w/v).

### Collection of human faecal material

Human faeces used to create the faecal emulsion was collected from healthy volunteers (N= 4, aged 18-75), without preceding antibiotic therapy (past three months). Faecal samples were transported in sealed anaerobic bags and placed in an anaerobic cabinet within 12 h of production. All samples were negative for the presence of CPE upon screening [(Biomerieux chrom ID® CARBA-SMART (carba- smart) & Cepheid Xpert® Carba-R assay (XCR)] and were pooled to produce ≈ 50 g of faeces. The subsequent pooled faecal material was emulsified with 500 mL of pre-reduced PBS and filtered through sterile muslin to create a smooth 10% w/v faecal slurry.

### CPE strains

Three clinical isolates of carbapenemase producing *K. pneumoniae* with distinct carbapenemase genes were used in this study. The carbapenemase genes selected were *bla* KPC (KPC) gene encoded on the PKpQiL-D2 plasmid, minimum inhibitory concentration (MIC) ertapenen (ERT) 4 mg/L; imipenem (IMI) 8 mg/L; meropenem (MER) 4mg/L, Oxacillinase-48 (OXA-48), MIC ERT 32mg/L; Mer 16mg/L, and New Delhi metallo-β-lactamase (NDM) MIC ERT 32mg/L; Mer 8mg/L; IMI 1mg/L, supplied and MICs tested by Public Health England (PHE), Leeds. Strains were reconstituted on blood agar from -80°C freezer strage, sub-cultured onto CPE selective media (Carba-Smart), and carbapenemase genes confirmed by PCR [Cepheid Xpert® Carba-R assay (XCR) multiplex real-time PCR assay and Check- Direct CPE Screen for BD MAX™ (CDCPE) multiplex real-time PCR assay].

### Experimental design

Three gut models were run in parallel for a total 38 days. The models were initially primed with pooled CPE negative faeces (from four healthy volunteers) and left for two weeks to allow the bacterial populations to equilibrate. Each model was exposed to increasing daily inocula of a different carbapenemase producing *K. pneumonia* strain (described above) from a 10-fold dilution series of a fresh overnight culture in nutrient broth (range 1.9-8.9 log_10_ cfu over 8 days) (Figure 1). points for this study. Period A represents the two week equilibration period following priming with faecal slurry. Period B involved the inoculation of CPE strains which began on day 15. Period C represents the post inoculation sampling period. The solid black dots represent sampling points.

**Fig 1.**
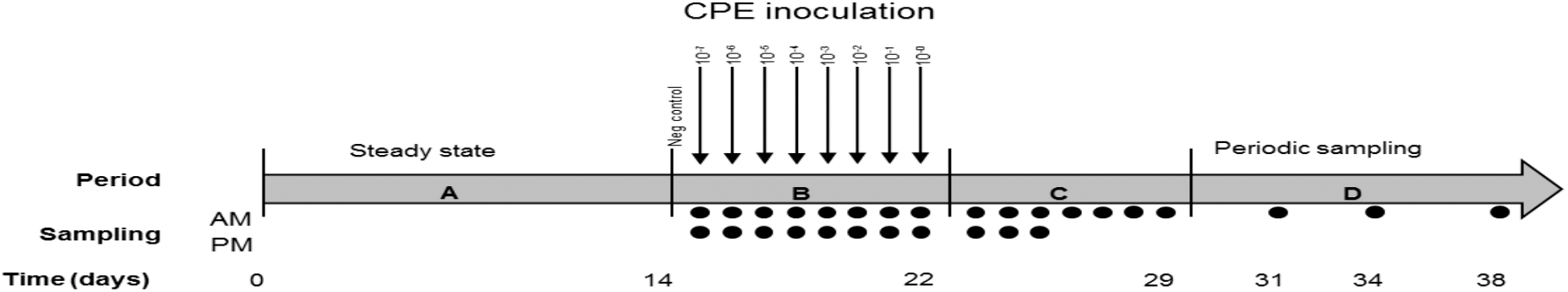
Schematic diagram illustrating the experimental timeframes and sampling

**Fig 1.**
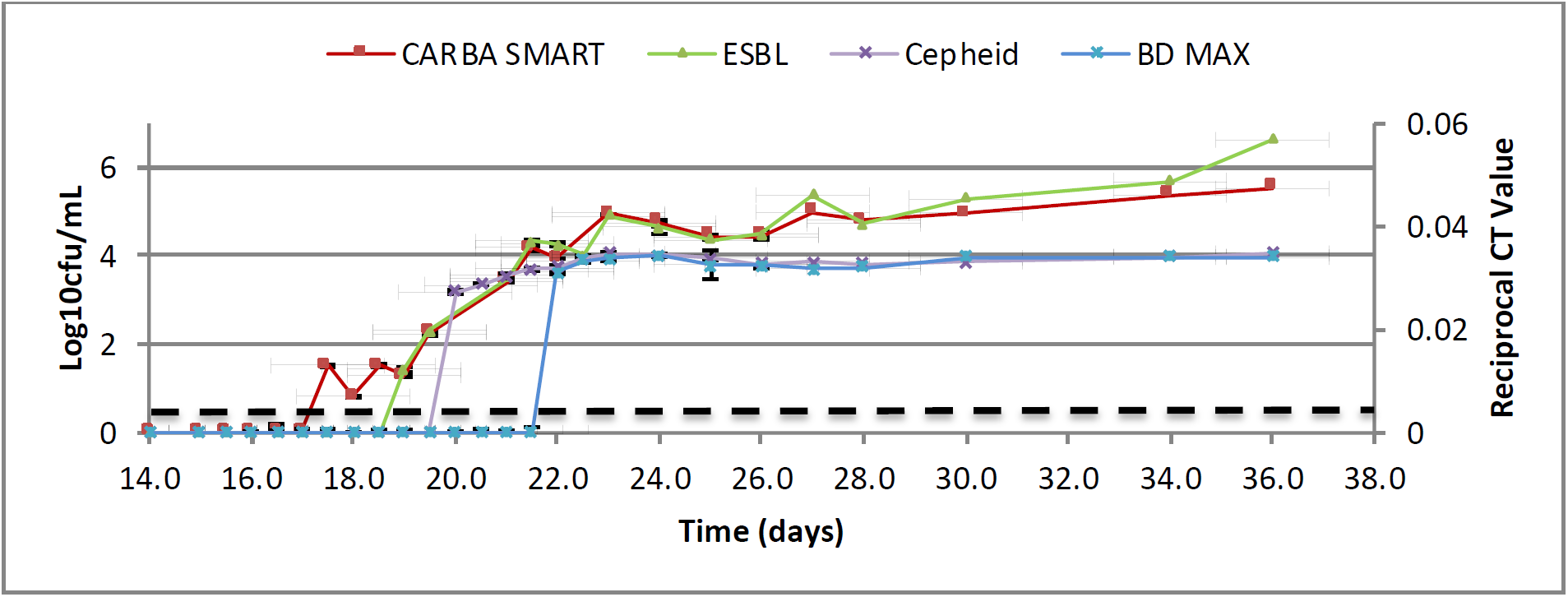
KPC Model: Comparison of detection limits between two CPE selective agars (mean og10 cfu/mL ± SE) and two molecular assays (mean 1/Ct ± SE) for the detection of KPC producing K. pneumoniae in Model A between periods B-D. The black dotted line represents the limit of detection for culture media (growth of a single colony; 0.82 log10 cfu/mL).

All three models ran as per Figure 1, with twice daily CPE screening of bacterial populations between days 15 and 25. CPE screening was then reduced to daily and subsequently twice weekly in further phases of the experiment

### Inoculation of Gut models with CPE isolates

Each model was inoculated with 1 m/L of an overnight culture (5 mL nutrient broth) of a different CPE strain (described previously) (Figure 1). Each strain was diluted in a 10-fold series to 10^-7^ in peptone water. The lowest (10^-7^) dilution was added to the model on the first day of inoculation (day 15). This was increased 10-fold daily until a neat solution was added on day 22 of the experiment. Cultures were enumerated on MacConkey agar to ensure comparable overnight cultures. This confirmed inoculum levels were as expected (Appendix 1).

### Population sampling

Indigenous gut microbiota populations were identified and enumerated daily using a variety of selective and non-selective agars, as previously described [20]. CPE populations were identified using five different detection tests which encompassed both selective agars and molecular methods. The selective agars used were BioMerieux chromID® ESBL (ESBL) and BioMerieux chromID® CARBA SMART (CARBA-SMART) and MacConkey agar containing 0.5 mg/L imipenem poured in-house (MAC-IMI). The molecular tests used to identify CPE genes were Cepheid Xpert® Carba-R assay (XCR) multiplex real-time PCR assay and Check-Direct CPE Screen for BD MAX™ (CDCPE) multiplex real-time PCR assay.

CPE populations were sampled (Figure 1). For culture based techniques approximately 5 mL of gut model culture fluid was extracted from vessel 3 (V3) of each model, with morning sampling for CPE populations occurring 1 h after CPE inoculation into the models (Figure 1, period B). The aliquots of gut model culture fluid were diluted in peptone water in a 10-fold dilution series. Samples were either diluted to 10^-3^ for CPE populations or up to 10^-7^ for enumeration of indigenous microbiota. Plates were inoculated in triplicate, using 50µl of each dilution series onto the CPE selective agars and 20 µL of the appropriate dilution for indigenous bacterial counts. Colonies were counted following 48 h incubation at 37°C, at an appropriate dilution to give estimation (log_10_ cfu/mL) of the CPE population within the model.

For the molecular platforms 50 µL of neat gut model fluid was assayed in triplicate, with the exception of using gut model fluid, rather than a rectal swab, this was in accordance with the manufacturer’s protocol. (Reference protocol here).Cycle threshold (Ct) values were recorded, along with the result (positive or negative) as determined by the manufacturer, using the threshold set for the assay. Reciprocal Ct values (1/Ct) and standard error (SE) were calculated.

### Data handling and analysis

Comparative analysis of all screening methods was undertaken using both single and strict criteria as described below:

Single criterion: 1. Any growth of Enterobacteriaceae on agar plates was considered positive (log_10_ cfu/mL reported); 2. For molecular assays, all reported Ct values were interpreted as positive up to _≤_40 cycles for XCR or _≤_50 cycles for CD CPE. A single positive result for any of the triplicate samples was recorded as positive (Table 2).

**Table 2.** Detection of CPE. During the twice daily sampling an evening sample is represented by adding 0.5 to the day. *^a^* Day positive for culture refers to the day the plate was inoculated not the day the plate was read, for molecular test refers to the day the test was completed. *^b^* Refer to methods, data handling and analysis.

| CPE gene | Screening method | Day positive <sup>a</sup><br>Single criterion<sup>b</sup> | Day positive <sup>a</sup><br>Strict criteria<sup>b</sup> |
| --- | --- | --- | --- |
| KPC Model | ESBL | 17.5 | 19.0 |
|  | CARBA-SMART | 17.5 | 17.5 |
|  | MAC-IMI | N/A | N/A |
|  | XCR | 20.0 | 20.0 |
|  | CDCPE | 19.5 | 22.0 |
| OXA 48 model | ESBL | 18.5 | 19.0 |
|  | CARBA-SMART | 17.5 | 18.0 |
|  | MAC-IMI | 17.5 | 19.0 |
|  | XCR | 18.0 | 20.5 |
|  | CDCPE | 18.5 | 19.5 |
| NDM Model | ESBL | 18.5 | 19.5 |
|  | CARBA-SMART | 18.5 | 19.0 |
|  | MAC-IMI | 19.5 | 19.5 |
|  | XCR | 19.0 | 20.0 |
|  | CDCPE | 17.0 | 21.0 |

Strict criteria of 1. Triplicate positive culture for selective media; 2. XCR: Using the internal algorithm reporting the sample as positive and samples were considered negative unless all three replicates were positive by the assay algorithm; 3 CD CPE: As there is no diagnostic algorithm associated with this assay, a clinical cut-off for a negative result of a Ct value _≥_35 was applied, as is often applied to routine diagnostic in-house PCR assays [21, 22] (Table 2). Again, samples were considered negative unless all three replicates were positive using this cut-off.

## Results

A total of 237 samples were tested using the five screening methods. No CPE strains were identified during the equilibrium phase (Figure 1.Period A) by either culture or molecular technique. For all assays, detection levels (as measured by log_10_ cfu/mL or 1/Ct) increased as the concentration of the inocula increased (Figure 2). Selective media detected CPE earlier or at the same time as molecular methods with the exception of MAC IMI, which showed poor performance overall; MAC IMI was subsequently discontinued in the KPC model (Figure 1).

**Fig 2.**
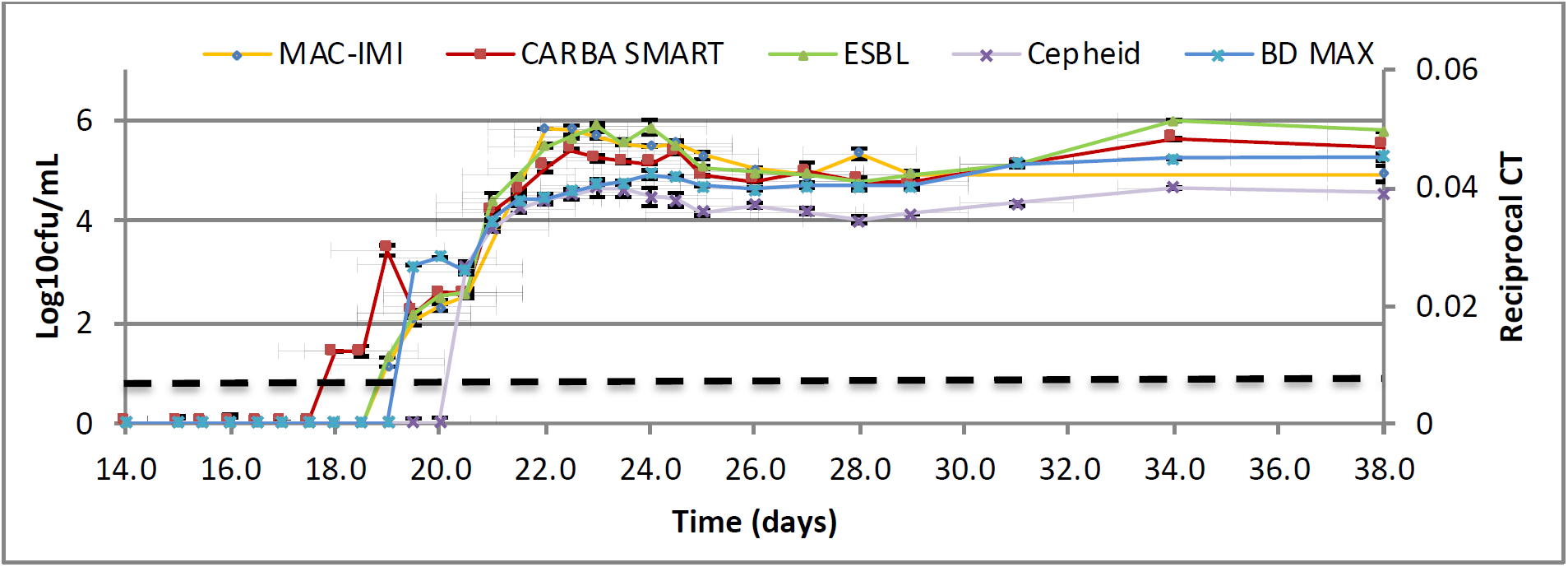
OXA 48 Model: Comparison of detection limits between two CPE selective agars (mean log10 cfu/mL ± SE) and two molecular assays (mean 1/Ct ± SE) for the detection of KPC producing K. pneumoniae in Model A between periods B - D. The black dotted line represents the limit of detection for culture media (growth of a single colony; 0.82 log10 cfu/mL).

### Detection of CPE within the gut model

CPE were detectable only after an inoculation of ∼4.9 log_10_ cfu/mL (Figure 1-3, Appendix Table 2). Populations increased with higher inocula, peaking at 5-6 log_10_ cfu/mL in vessel 3 for all gut models irrespective of CPE strain (Figure 1-3). Interestingly, CPE populations remained stable within the gut microbiota on cessation of CPE inoculation for all models. Following the final inoculum of ∼8.9 log_10_ cfu/mL, carbapenem producing (CP) *K. pneumoniae* populations stabilised at 3- 5 log_10_ cfu/mL for the remainder of the experiment, with no evidence of ‘washing out’ of CPE populations in any of the three models.

**Fig 3.**
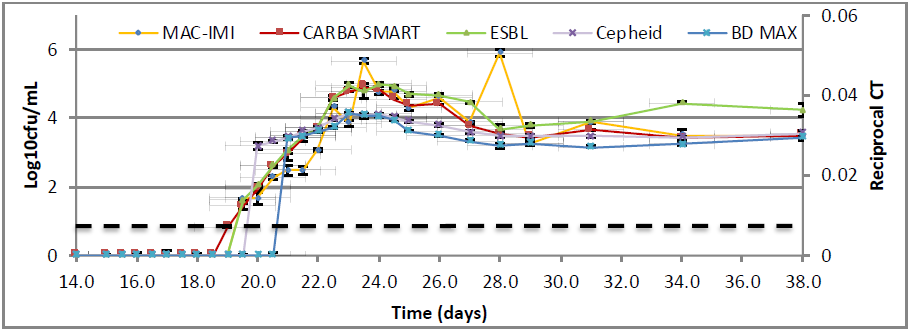
NDM model: Comparison of detection limits between two CPE selective agars (mean log10 cfu/mL ± SE) and two molecular assays (mean 1/Ct ± SE) for the detection of KPC producing K. pneumoniae in Model A between periods B - D. The black dotted line represents the limit of detection for culture media (growth of a single colony; 0.82 log10 cfu/mL).

Ct values stabilised in all models (Figure 1-3), and have been reported as 1/Ct to allow graphical presentation with viable counts. Correlation of Ct Values and viable counts showed that at higher culture density there were corresponding lower Ct values. For both XCR and CD CPE, a Ct valve of ≤35 corresponded to triplicate positive culture and hence was predictive of colonisation.

### Comparative evaluation of CPE screening methods

Performance of screening methods was reported in days from the onset of the experiment when the assay became positive (as per data handing and analysis described above) (Table 2). The MAC IMI data are incomplete as this agar was technically unpredictable and diagnostically difficult to count; consequently, it was discontinued. The Carba-Smart agar detected KPA and OXA 48 first using both simple and strict criteria. For NDM, CD CPE detected this strain first under the simple criteria; however, use of strict criteria showed that Carba-Smart identified this strain first. The XCR detected CPE later than other methods for KPC and OXA48 strains. The XCR assay detected all enzymes later than the CD CPE assay; however, following application of strict criteria, XCR detected KPC and NDM before CD CPE (Table 2).

### Relative sensitivity of CPE screening methods

The MAC-IMI agar displayed a lower limit of detection (LOD) of 1.66 log_10_ cfu/ml. Both commercial chromogenic agars had similar sensitivity and a lower LOD of 0.82 log_10_ cfu/mL. The relative sensitivity and specificity of the molecular methods were calculated compared to the reference method, which was triplicate positive culture (Carba-Smart) and summarised in Table 2. A molecular test was considered positive if there was detection of at least one CP gene, regardless of the internal algorithm or CT value. XCR showed decreased sensitivity but increased specificity compared with CD CPE.

**Table 2.** Sensitivity & specificity of molecular tests in reference to culture with 95% confidence interval.

| Method | Gene | Sensitivity % | Specificity % |
| --- | --- | --- | --- |
| XCR | KPC | 83.33 (73.62-90.58) | 100 (97.62-100) |
|  | OXA 48 | 87.30 (76.50-94.35) | 99.43(96.84- 99.99) |
|  | NDM | 96.49 (87.89-99.57) | 99.44 (96.94-99.99) |
| CD CPE | KPC | 97.62 (91.66-99.71) | 98.04 (94.38-99.59) |
|  | OXA | 92.06 (82.44-97.37) | 87.14 (82.14-92.37) |
|  | NDM | 98.25 (90.61-99.96) | 86.67 (80.81-91.27) |

### Detection of strains not inoculated into the models

Any detection of a CPE gene that has not been directly inoculated into the model was considered a false positive result (Table 3). CD CPE had a higher rate of false positives compared with XCR, particularly with the detection of the OXA 48 and NDM genes.

**Table 3.** The number of false positive result CPE gene detection results by each molecular test.

| Method | KPC | OXA48 | NDM | IMP1 | VIM |
| --- | --- | --- | --- | --- | --- |
| XCR | 0 | 1 | 0 | 0 | 0 |
| CD CPE | 3 | 19 | 15 | 0 | 0 |

## Discussion

Using a simulated human colonic environment, we have demonstrated CPE colonisation with three distinct carbapenemase producing strains, suggesting this is a generic ability of CPE isolates rather than a unique capacity of individual strains. These colonisation events occurred without antibiotic selective pressure; thus, we have shown that a dysbiotic flora is not a prerequisite for CPE colonisation. The highest CPE viable counts (5.4 log10 cfu/ml) at the end of the experiment (day 38) was seen in the OXA model, reflecting recent data showing that this a rapid coloniser [23, 24]; however, counts of KPC (4.6 log10 cfu/ml) and NDM (3.9 log10 cfu/ml) also remained relatively high. Our selected OXA strain had a relatively high MIC of both ERT and MER, which might influence biomass as well the sensitivity of the selective agar for its detection.

Screening for faecal colonisation with CPE is routine clinical practice worldwide and is advocated by multiple health agencies including Public Health England (PHE, 2013, Canton et al., 2012). Screening algorithms used for CPE detection differ greatly between institutions, based on infrastructure, cost and prevalence, although the aim of CPE detection at low cost, high accuracy and high throughput remains the same. Analysis of screening performance demonstrated that the Carba-Smart selective media had the highest sensitivity for CPE detection of all methods investigated. Carba-Smart has been previously shown to be superior to other selective media [25] when tested using rectal swabs.

We found the ESBL agar was less sensitive than the Carba-Smart; however, it did detect the presence of CPE sooner than the molecular platforms (XCR, CD CPE). The poured in house MAC-IMI posed significant technical problems with plating, enumeration and reading of the individual colonies and non-reproducible results. The instability of IMI in media (Turgeon and Desrochers, 1985) has been demonstrated and was likely responsible for the poor performance we observed. Vrioni et al., 2012 used plates poured in house incorporating MER, which proved more stable.

Given the performance of the selective media, we calculated the sensitivity and specificity of the PCR based tests against the standard of triplicate positive culture (Carba-Smart). It should be noted that the methodology used here cannot measure the specificity of the agars. Also, we did not investigate for isolates with ESBL or Amp C/ porin mutations, which may have the phenotypic appearances of CPE. XCR showed decreased sensitivity but increased specificity for KPC and OXA48 compared with CD CPE. Both methods had similar sensitivity for NDM, but XCR had higher specificity. It is worth noting that the PCR based assays are intended to be used with formed stool or rectal swab rather than gut model fluid, and so this may have impacted performance. To minimise comparative bias we used 50 µL of gut model fluid as both the media and molecular inoculum. However, the use of 50 µL of gut model fluid rather than directly plating a rectal swab resulted in the use of a higher inoculum, which may have increased the sensitivity of the agar method.

Lau et al., 2015 compared the sensitivity between CD CPE and selective media [HardyCHROM CRE agar (Hardy Diagnostics)] using two rectal swabs, one diluted into saline for plating (inoculum 2 drops) and a second that was mixed with lysis buffer and used as PCR template (50µL aliquot). They showed culture was at least as sensitive if not better then CD CPE in a LOD analysis. Similarly, they showed a Ct value of ≤35 correlated with a positive culture result, further corroborating our findings. Tato et al., 2016 conducted a multisite investigation of XCR compared with culture (MacConkey broth containing 1 mg/L of MER and subcultured to a MacConkey agar plate with a 10-μg MER disk) and DNA sequencing. Sensitivity and specificity were calculated as 96.6% and 98.6%, respectively [26], which was a higher sensitivity but similar specificity to our study. Individually, VIM had the lowest sensitivity of 93.5%, and NDM had the highest (100%), while KPC had a sensitivity of 96.7%, which was considerably higher than we recorded (83.3%).

Culture allows detection of non-targeted carbapenemase genes, and also has the capacity to detect non carbapenemase producing carbapenem resistant strains, in contrast to PCR based techniques, which will only detect a finite set of pre-programmed targeted resistance genes. However, the use of traditional phenotypic methods does have inherent drawbacks. Firstly, the data presented in table 2 represent the day the plate was inoculated rather than enumerated (to reflect inocula at that time); hence, in clinical reporting times there would be a 24 h lag (at least) required for incubation. The use of molecular tests negates this incubation time. The Public Health England toolkit (PHE, 2013) suggests pre-emptive source isolation of any high risk individuals until a negative result is available, and so such a lag period may not have a deleterious effect on patient outcomes. Secondly, selective media do not provide information on the genetic basis of the resistance, unlike the real time PCR assays that identify the most common CPE genes. Knowledge of the specific carbapenemase genes is crucial in resistance surveillance, and is of increasing clinical value as application of carbapenamase enzymology to treatment strategies [27] is investigated.

The CD CPE showed a higher sensitivity for CPE detection but also produced a higher rate of discrepant results, 15% (n=37/237) versus 0.4% (n=1/237), when compared with XCR. These possible false positive results were primarily centred on OXA 48 and NDM resistance genes. It is possible that these false positives were true positives, although this is unlikely as the results were sporadic and not reproducible. The CD CPE assay is a multistep open chamber system and therefore contamination could potentially account for the false positive rate; however, our false positive rate of 15% corroborates previous studies [22, 28, 29]. This high false positive rate undermines the use of this assay for infection control practices.

## Conclusion

The in-vitro human gut model is a novel way to evaluate CPE screening performance, notably allowing direct comparison of a variety of screening platforms for a range of test organisms in a controlled environment. Using a simulated human colonic environment, we have shown that CPE exposure can lead to insertion, multiplication and persistence of CPE strains within the human microbiota, providing evidence of new colonisation events. We have shown insight into the performance of five CPE recovery methods, and identified Carba-Smart as the most sensitive method. Chromogenic agars provide convenient and inexpensive CPE screening tools, albeit yielding slower results without enzymatic information.

## Appendix 1

**Table A1.** Approximate numbers of CPE added to the model

| Day | Dilution | Approximate inocula |
| --- | --- | --- |
| 15 | 10<sup>-7</sup> | 1.90 log<sub>10</sub> cfu/mL |
| 16 | 10<sup>-6</sup> | 2.90 log<sub>10</sub> cfu/mL |
| 17 | 10<sup>-5</sup> | 3.90 log<sub>10</sub> cfu/mL |
| 18 | 10<sup>-4</sup> | 4.90 log<sub>10</sub> cfu/mL |
| 19 | 10<sup>-3</sup> | 5.90 log<sub>10</sub> cfu/mL |
| 20 | 10<sup>-2</sup> | 6.90 log<sub>10</sub> cfu/mL |
| 21 | 10<sup>-1</sup> | 7.90 log<sub>10</sub> cfu/mL |
| 22 | neat | 8.90 log<sub>10</sub> cfu/mL |

## Appendix 2

**Figure A2:**

Appendix 2.
| Agar | Target population |
| --- | --- |
| Nutrient agar | Total facultative anaerobes |
| MacConkey Agar No3 | Lactose fermenting Enterobacteriaceae |
| Kanamycin Aesculin Azide Agar | Enterococcus spp. |
| Fastidious Anaerobe Agar | Total anaerobes (Faculative + Obligate) |
| Bacteroides Bile Esculin | Bacteroides fragilis group |
| LAMVAB | Lactobacillus spp |
| Beerens Agar | Bifidobacterium spp. |

